## Supplementary Material for "Deep phenotypic profiling of neuroactive drugs in larval zebrafish"

### Supplementary Material for
### Deep phenotypic profiling of neuroactive drugs in larval zebrafish

**Leo Gendelev[1], Jack Taylor[1,5], Douglas Myers-Turnbull[1], Steven Chen[2], Matthew N. McCarroll[1,2], Michelle R. Arkin[2], David Kokel[1 *], Michael J. Keiser[1,2,3,4 *]**

1. Institute for Neurodegenerative Diseases, University of California, San Francisco, San Francisco, CA, USA
2. Department of Pharmaceutical Chemistry, University of California, San Francisco, San Francisco, CA, USA
3. Department of Bioengineering and Therapeutic Sciences, University of California, San Francisco, CA, USA
4. Bakar Computational Health Sciences Institute, University of California, San Francisco, San Francisco, CA, USA
5. UCSF Weill Institute for Neurosciences Memory and Aging Center, University of California, San Francisco, CA, USA

### Supplementary Figure Legends

#### Figure S1. Random forest classification examples

Examples of using the random forest classifier to bin NT-650 drugs into three categories: active, inactive, and toxic. (a) Toxic example (Butaclamol). 2 of the 7 traces have a toxic probability > 0.8 (red lines), so we classify this drug as toxic. (b) Inactive example (2-MPDQ). 3 of the 7 traces have an inactive or "DMSO" probability > 0.5 (grey lines), so we classify this drug as inactive. (c) Active examples (Yohimbine). 6 of the 7 traces have a drug probability > 0.5 (green lines), so we classify this drug as active.

#### Figure S2. Model soundness checks

We perform a set of soundness checks for the fully randomized screen. (a) We train the Twin-NN model with completely randomized input features (motion index time series) and (b) with randomized output labels. In both cases, the result is a model that cannot distinguish between positive and negative pairs. (c) We train Twin-NN models where positive and negative pairs are defined by plate distance within a neighbor-cutoff of 2.0 and (d) with a neighbor-cutoff of 5.0. If a pair is within the neighbor-cutoff distance, it is a positive pair and negative otherwise. In both cases, these models cannot distinguish positive from negative pairs.

#### Figure S3. Correlation distance behaviorome UMAP

Same as Figure 4 (Results), but using correlation distance.

#### Figure S4. Correlation distance vs Twin-NN "phenosearch" top-ranked control-well counts

We plot the number of control (DMSO) wells (y-axis) in the pheno-search results for the NT-650 compounds compared with the strength of the phenotype (rf_score, x-axis) (a) using Twin-NN distance for the phenosearch (b) using correlation distance. Strikingly, there is a highly populated area of points with Twin-NN, average-strength rf_score (0.2-0.5) with few control wells, whereas very few points of this type with correlation distance. This suggests Twin-NN distance is more effective at finding drug-like novel compounds in pheno-searches than correlation distance, especially for compounds with weaker or average-strength phenotypes.

#### Figure S5 – S13. Concentration-response competition binding curves of selected compounds at selected receptors.

Results (mean ± SEM) from a minimum of 3 independent assays (each in triplicate) were normalized, pooled, and analyzed using the built-in competition binding function in the GraphPad Prism V10.

### Figure S1: Random forest classification examples

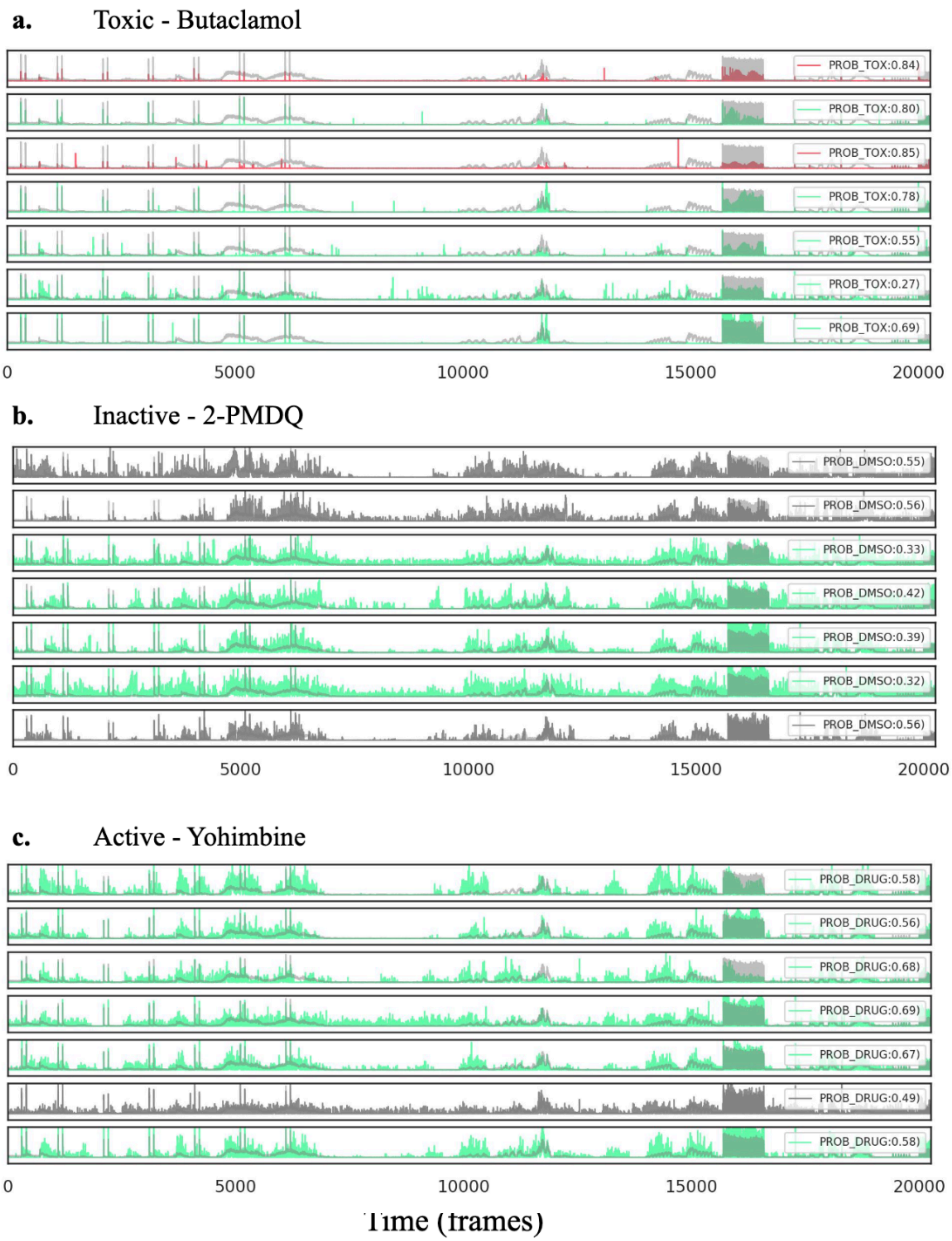

### Figure S2. Model soundness checks

Random Features (MI Values)

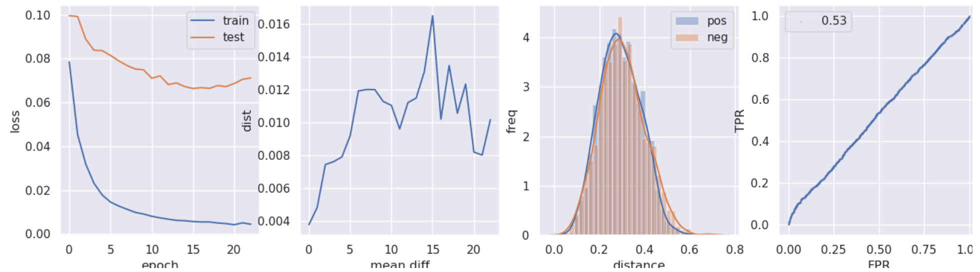


Randomized Labels

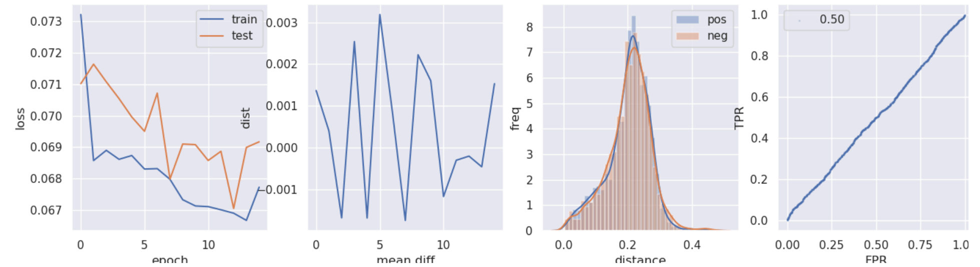


Predicting well dist (neighbor cutoff 5.2)

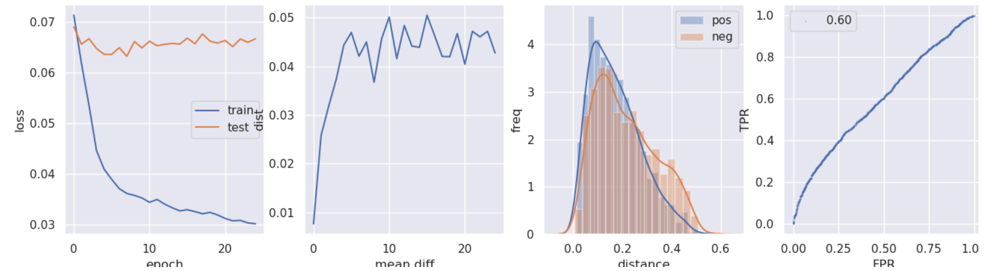


Predicting well dist (neighbor cutoff 2.0)

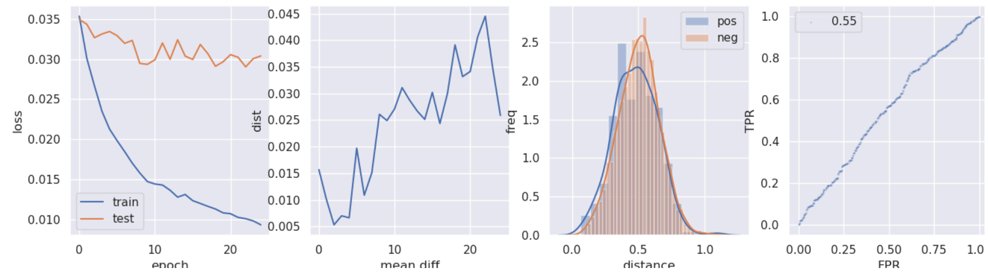

### Figure S3. Correlation distance behaviorome UMAP

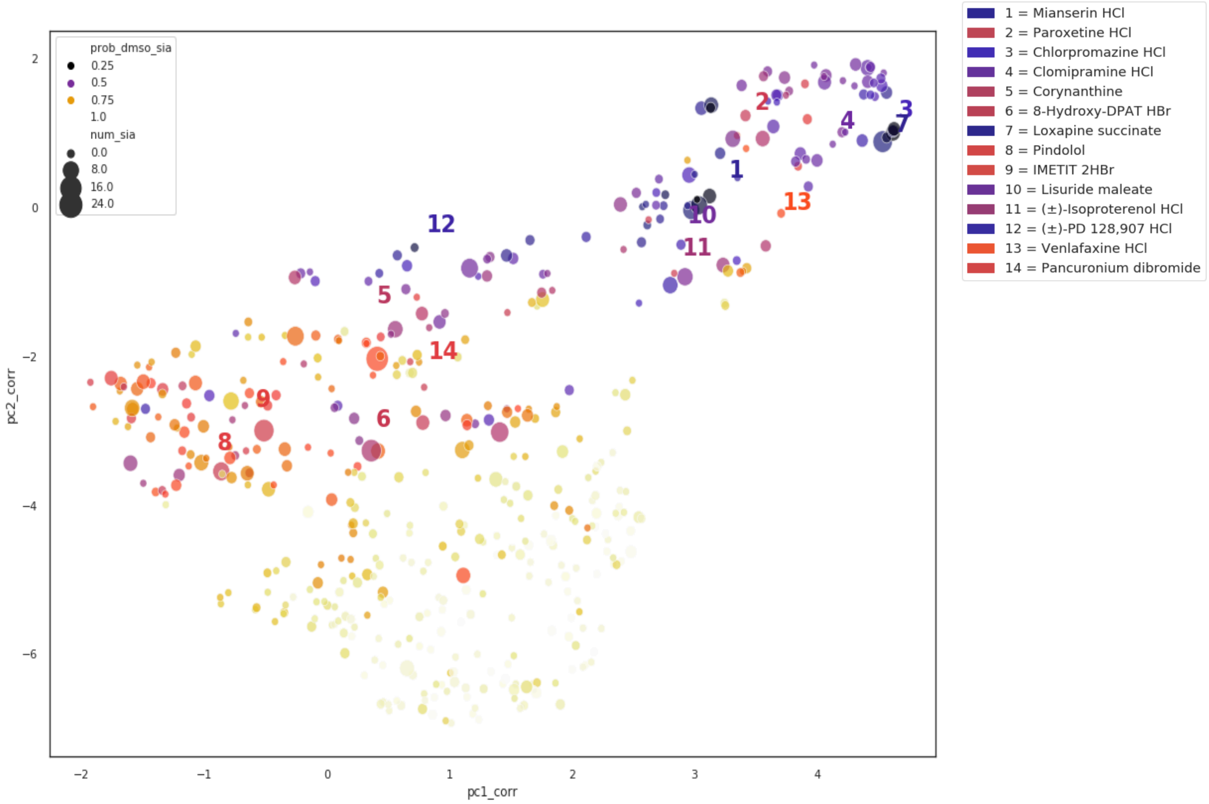

#### Figure S4. Correlation distance vs Twin-NN "phenosearch" top-ranked control-well counts

a.

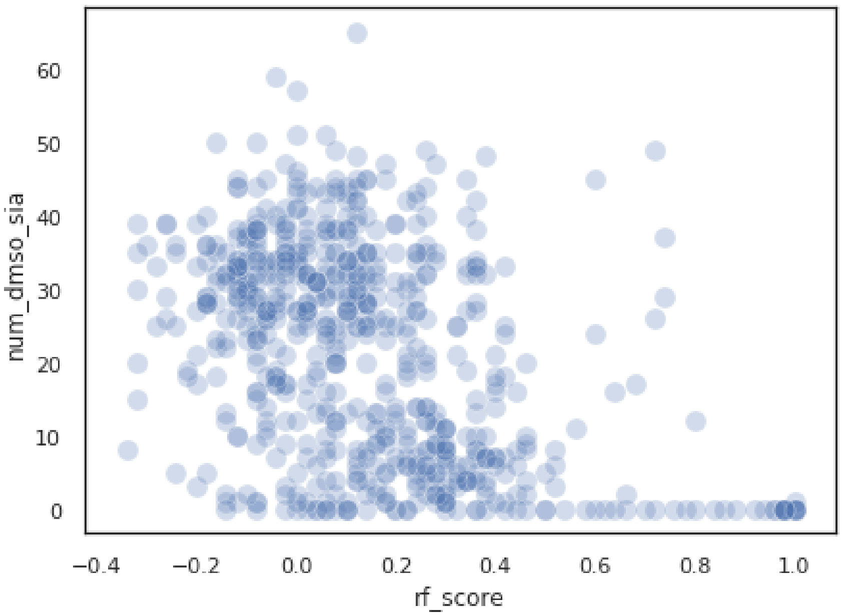


b.

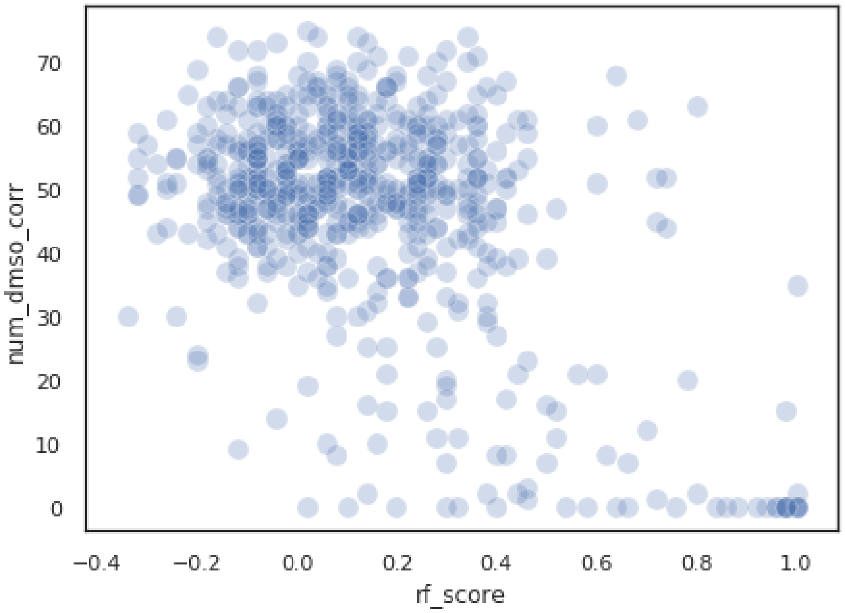

**Figure S5. 5-HT1A Secondary Binding Curves**

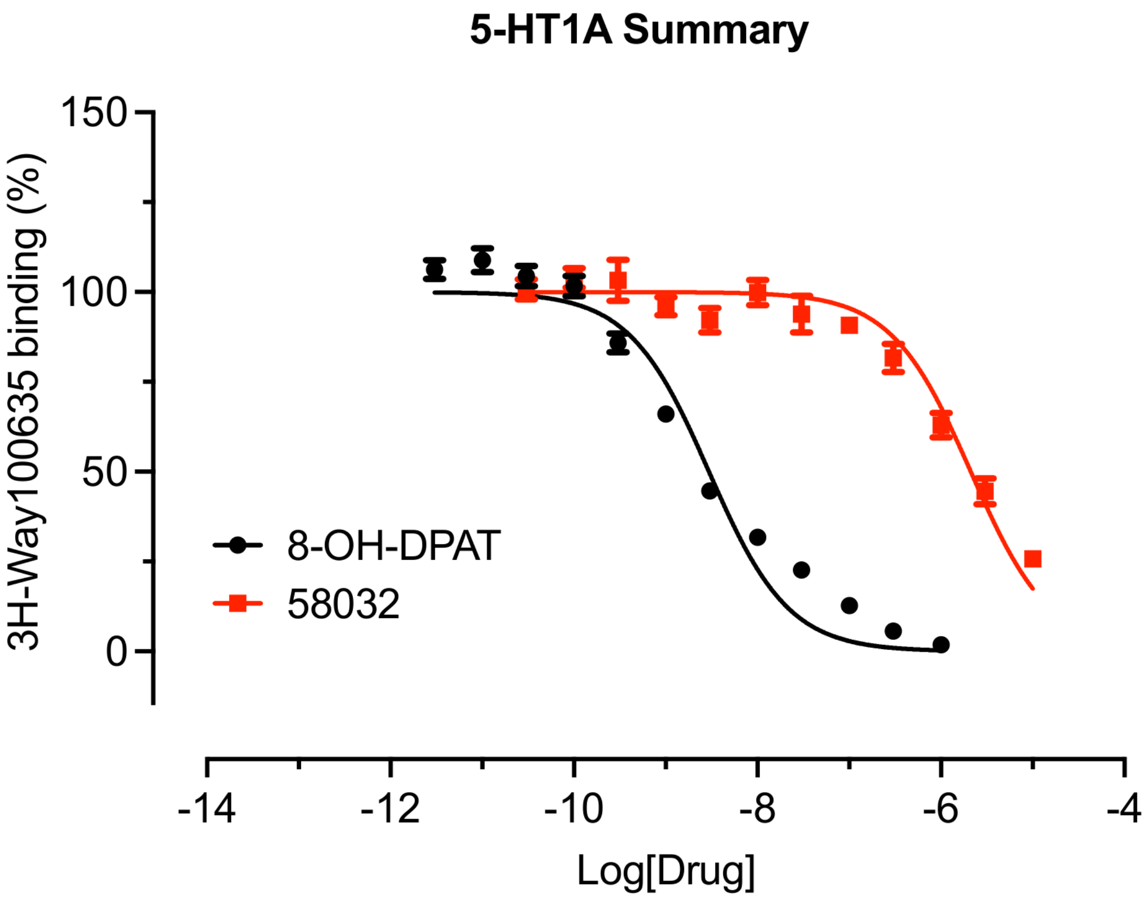

**Figure S6. 5-HT2A Secondary Binding Curves**

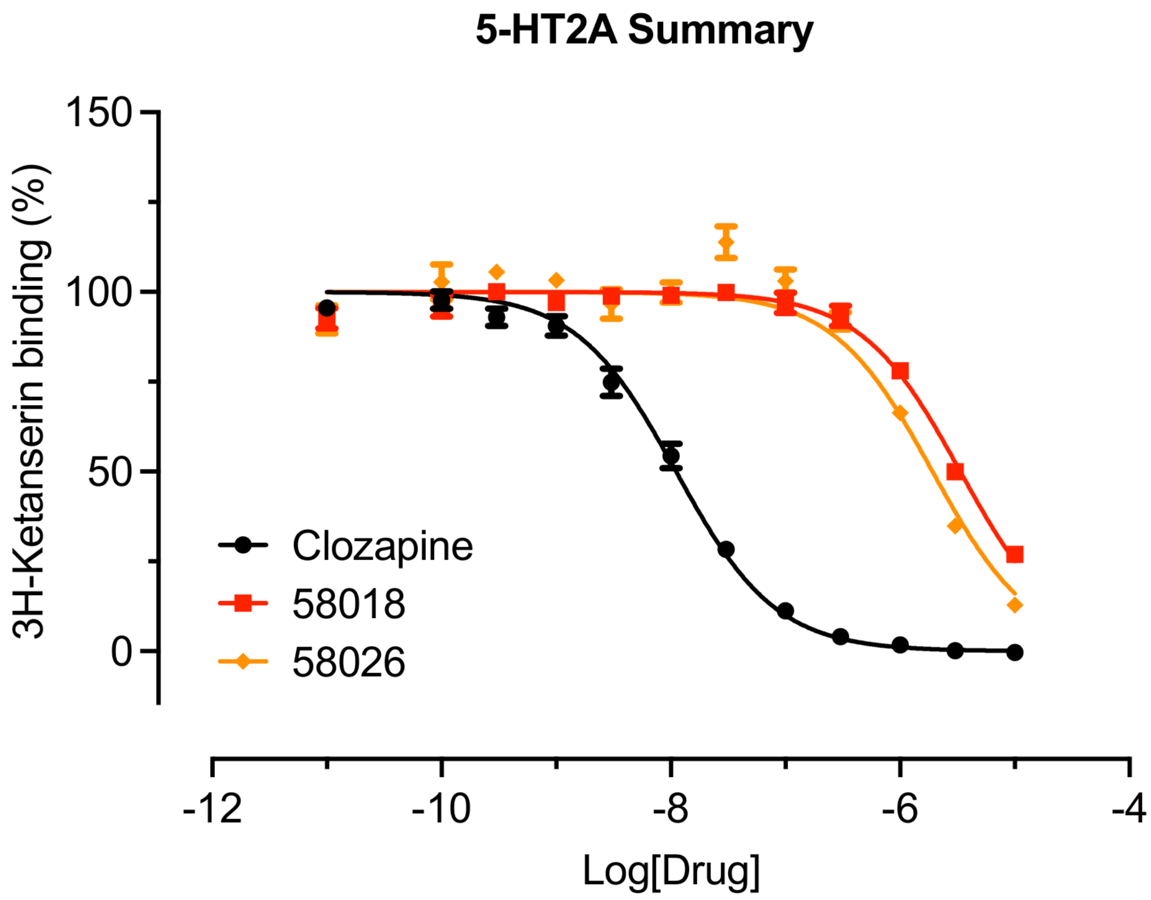

**Figure S7. 5-HT2B Secondary Binding Curves**

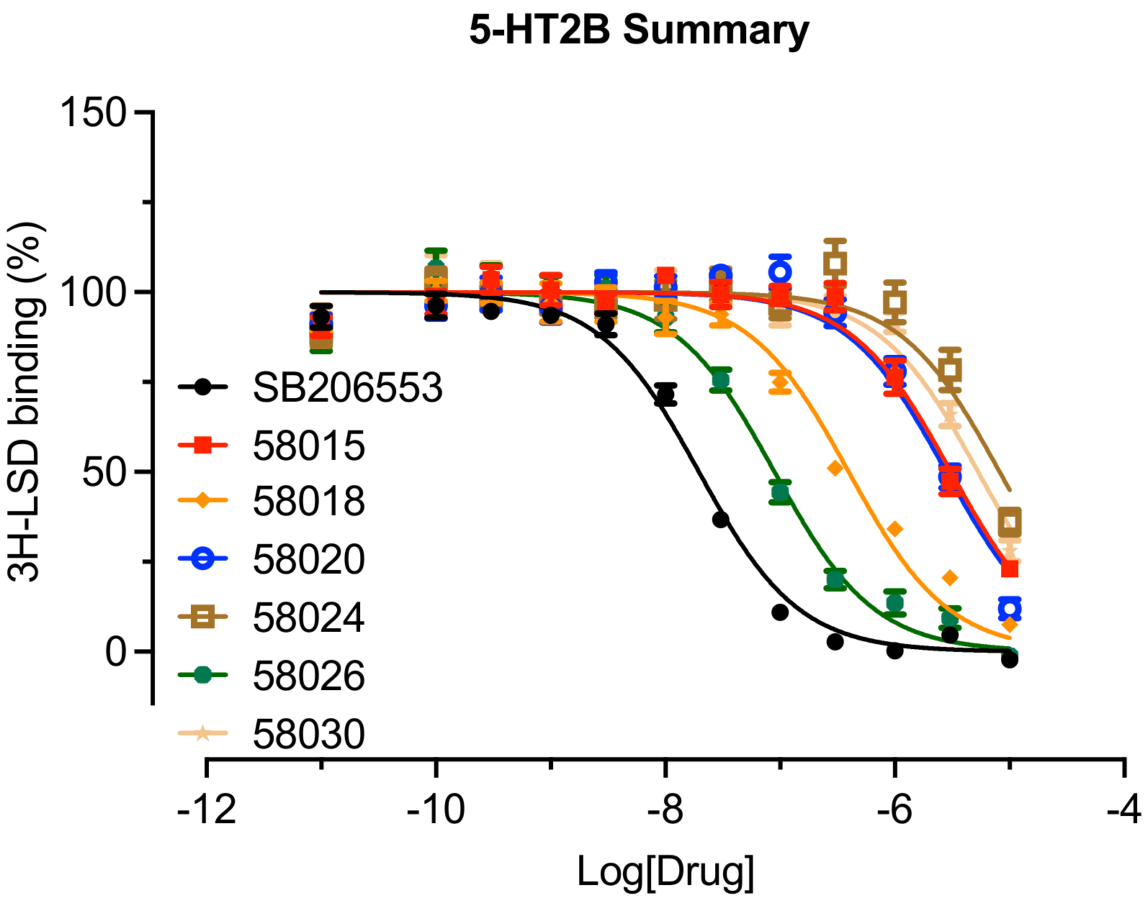

**Figure S8. 5-HT2C Secondary Binding Curves**

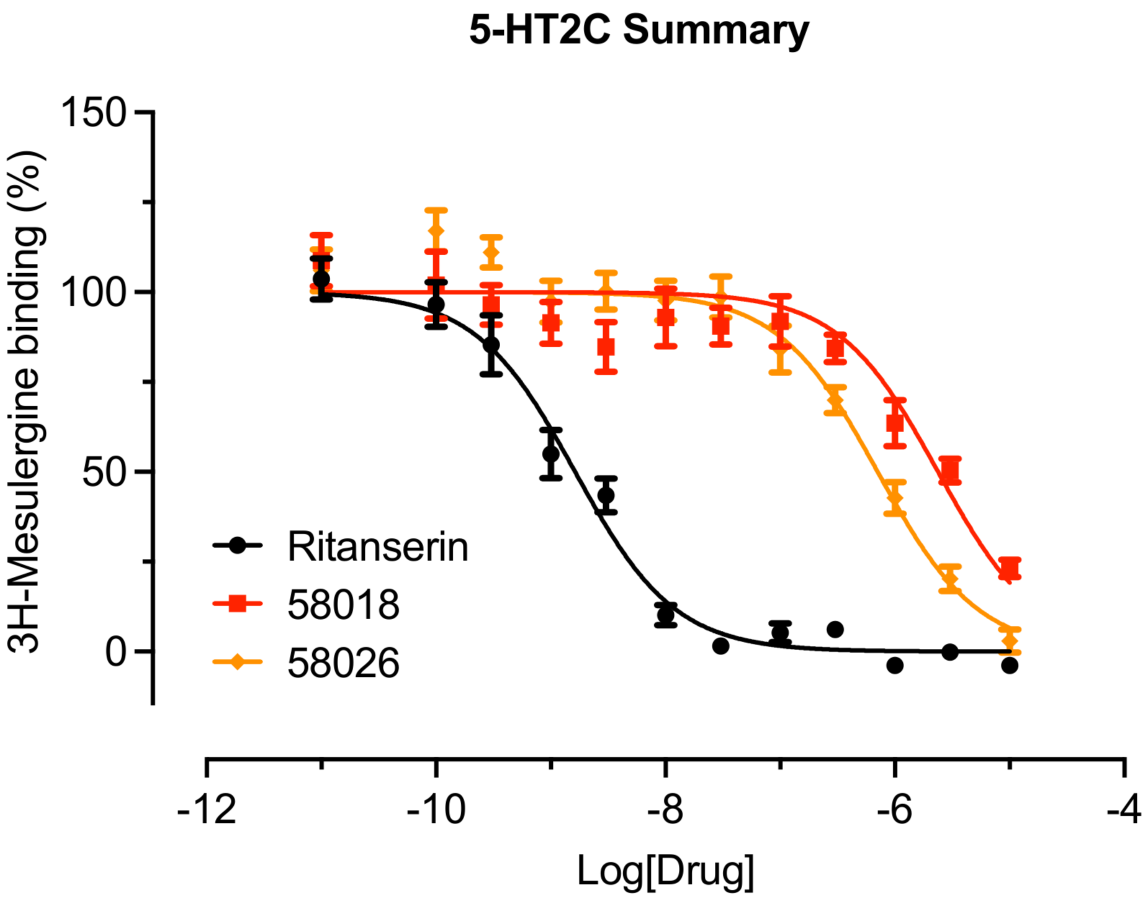

**Figure S9. 5-HT7A Secondary Binding Curves**

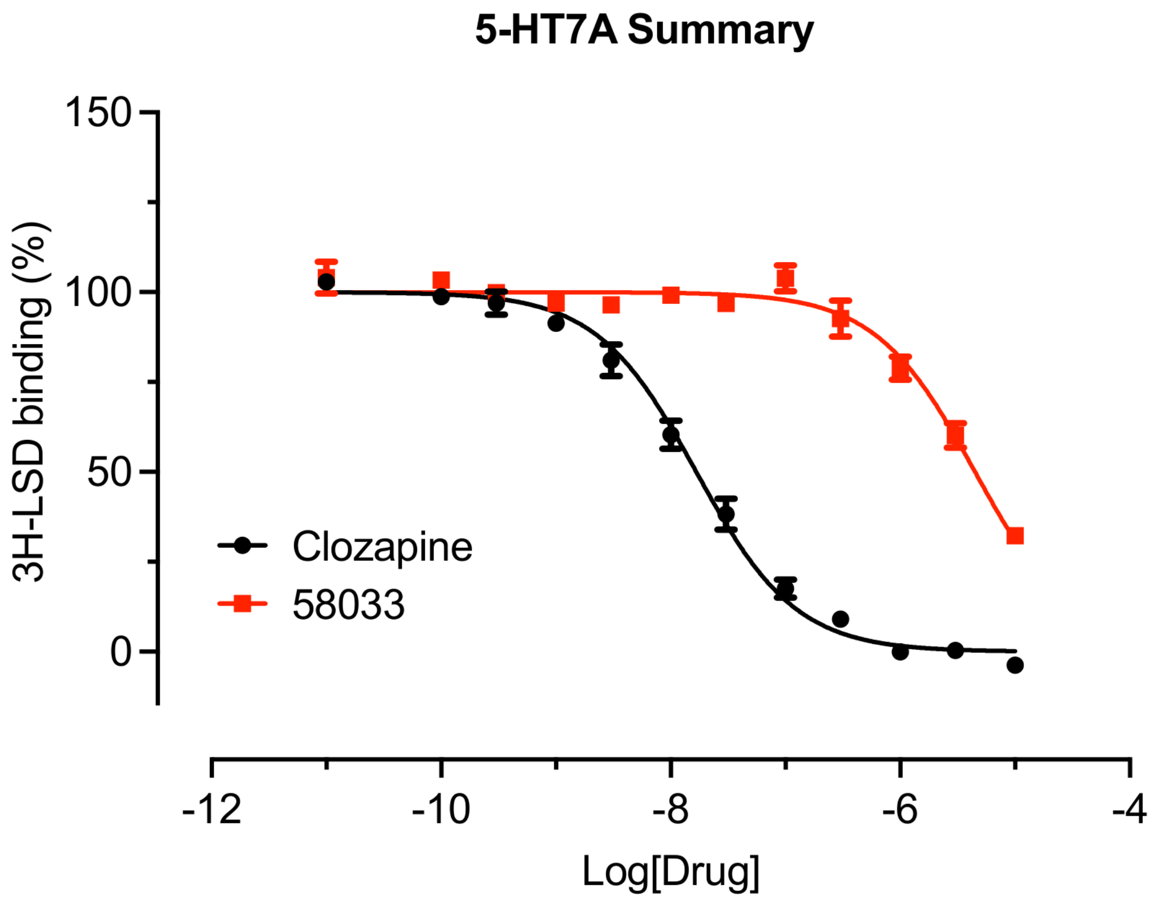

**Figure S10. Dopamine D2 Receptor Secondary Binding Curves**

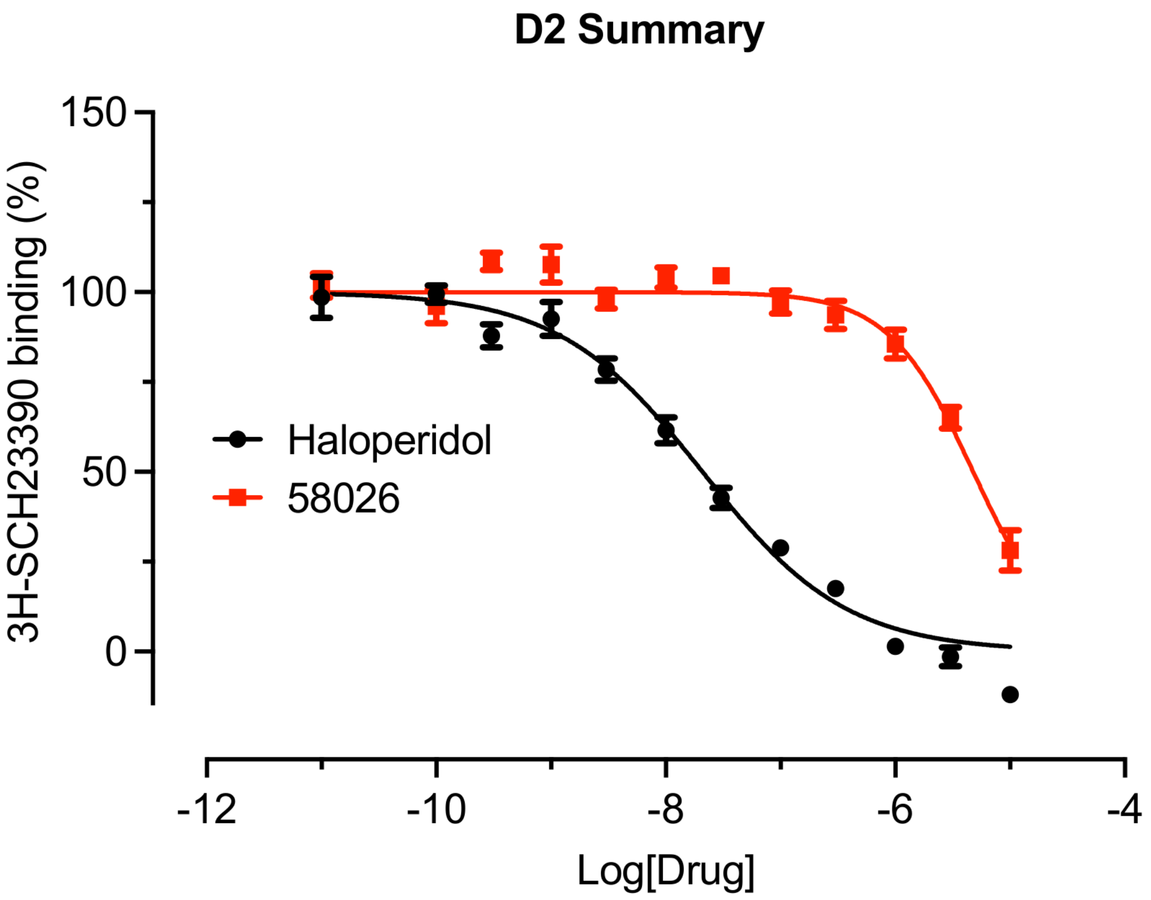

**Figure S11. Histamine H3 Receptor Secondary Binding Curves**

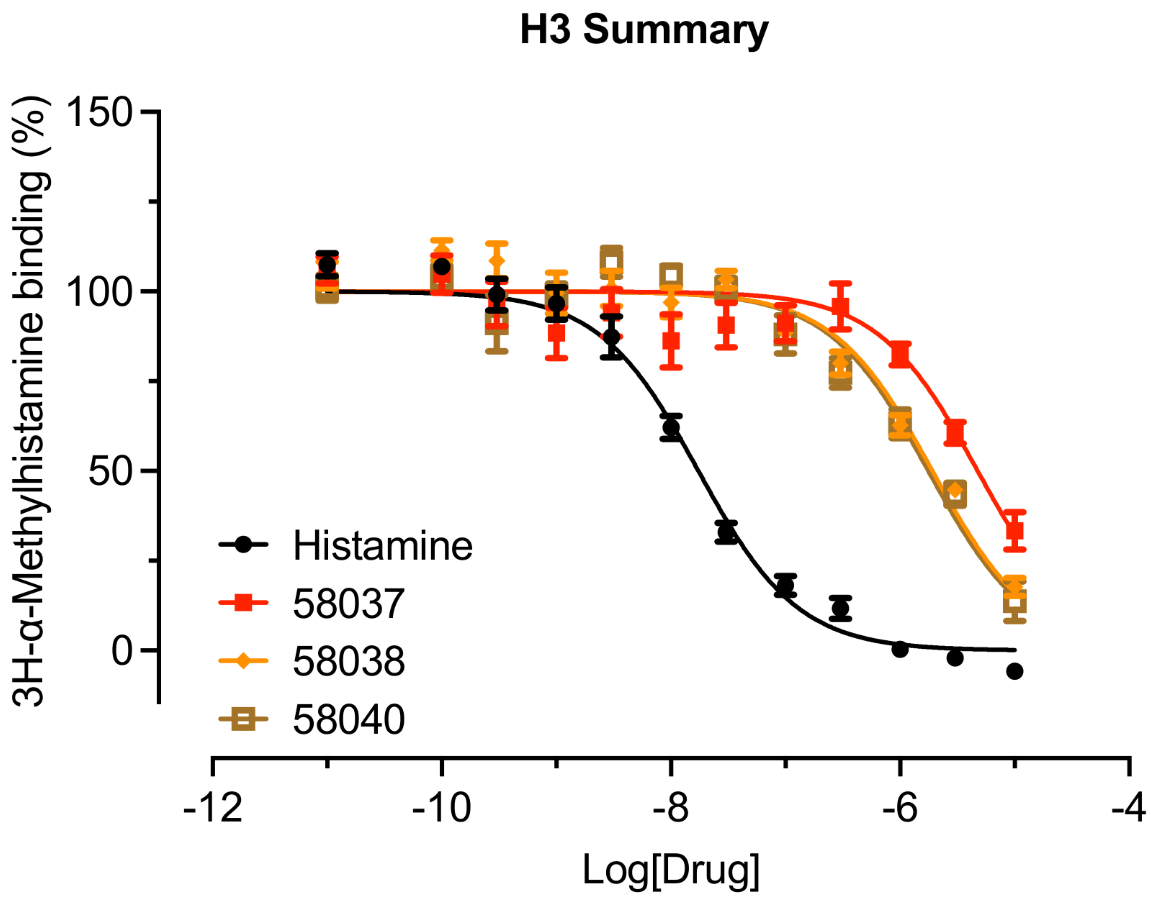

**Figure S12. Histamine H4 Receptor Secondary Binding Curves**

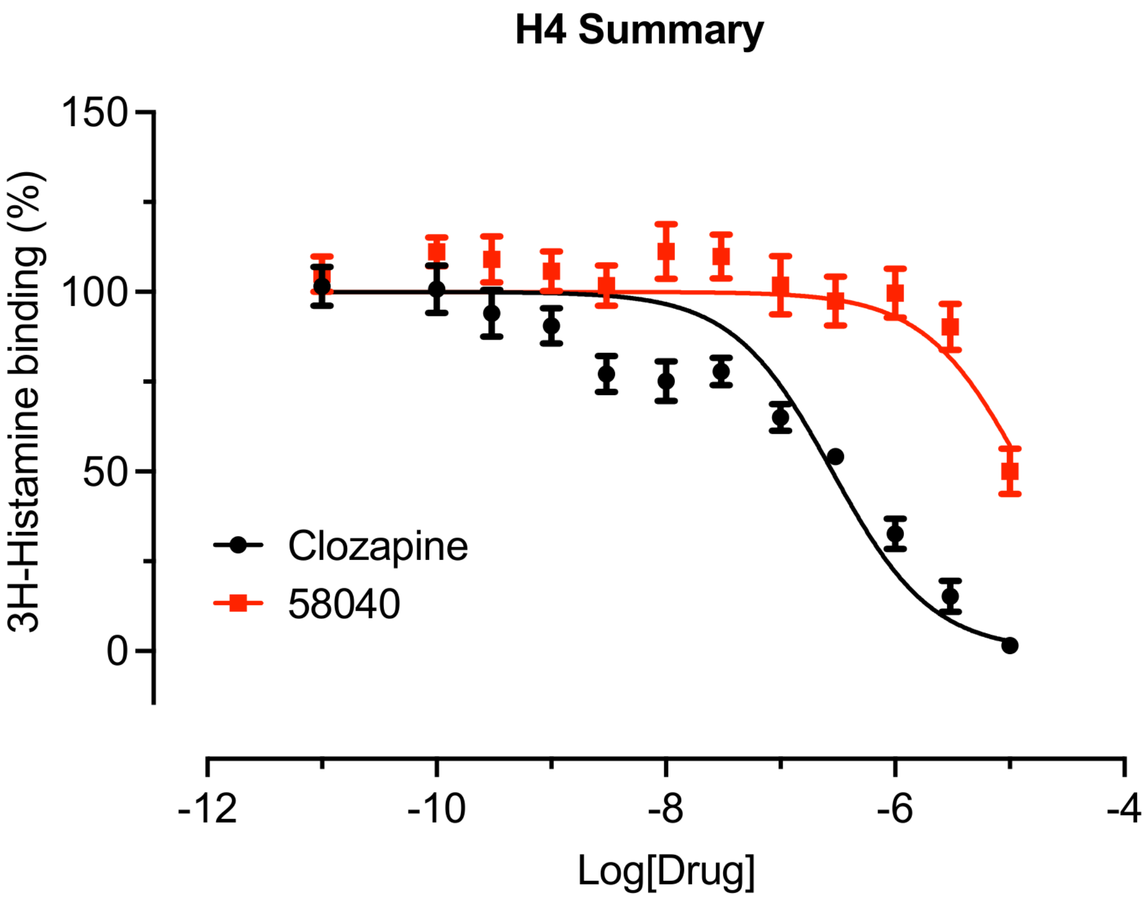

**Figure S13. Alpha 2C Adrenergic Receptor Secondary Binding Curves**

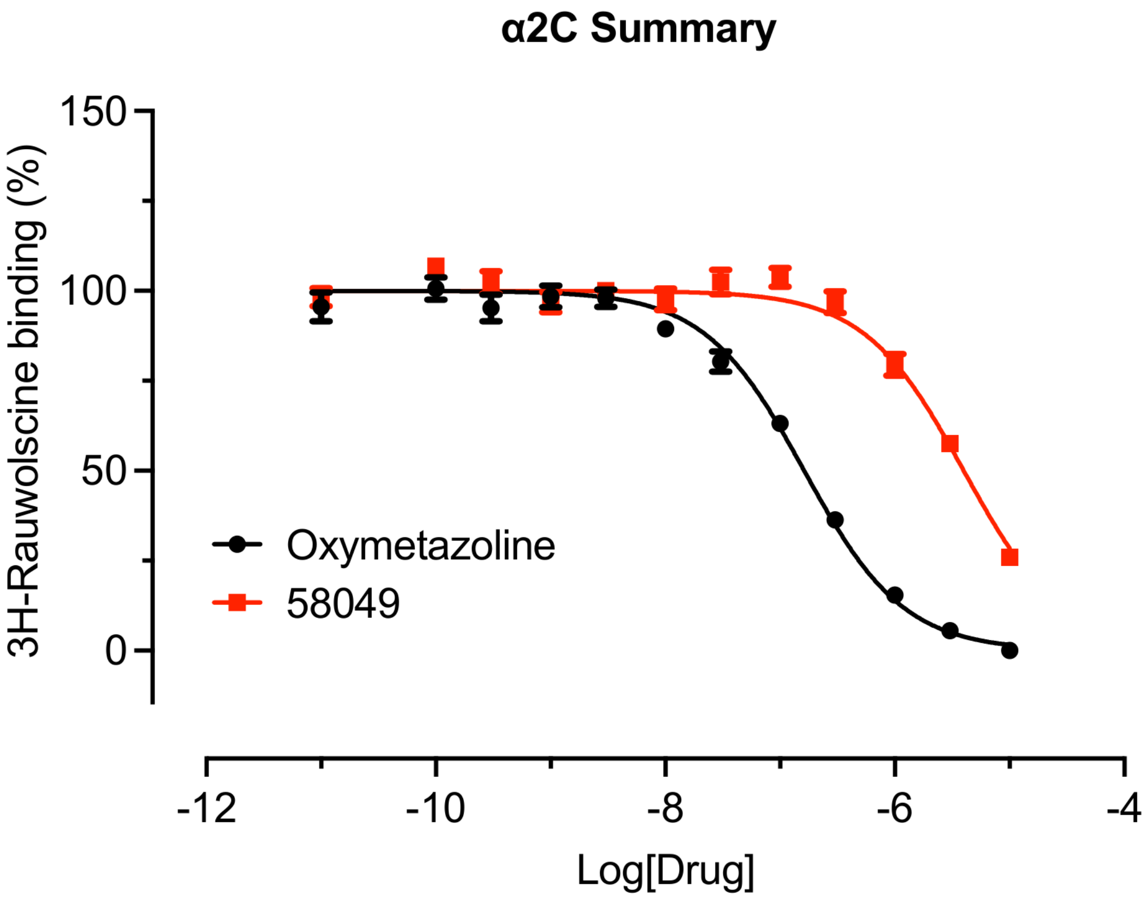

### Table T1: Internal ID to Hit2Lead compound ID mapping

| Internal_ID | Hit2Lead_ID |
|---|---|
| c29237 | 9210449 |
| c26692 | 9239432 |
| c29296 | 9212284 |
| c25328 | 9207145 |
| c26732 | 9240531 |
| c33446 | 9302621 |
| c33448 | 9306584 |
| c29676 | 9216089 |
| c33858 | 9336481 |
| c29440 | 6423322 |
| c32656 | 9275801 |
| c26429 | 9232516 |
| c33965 | 9343970 |
| c33151 | 9284190 |
| c31211 | 9247328 |
| c25518 | 9211046 |
| c27884 | 9265565 |
| c25477 | 9209700 |
| c24594 | 4033665 |
| c31386 | 9251970 |
| c34037 | 9201123 |
| c33553 | 9308643 |
| c33255 | 9287933 |
| c31508 | 9254844 |
| c31447 | 9253440 |
| c28263 | 9277938 |
| c33507 | 9309842 |
| c26398 | 9232267 |
| c26521 | 9232718 |

| | |
|---|---|
| c33914 | 9340973 |
| c30977 | 9243029 |
| c28263 | 9277938 |
| c27284 | 9253599 |
| c33507 | 9309842 |
| c26398 | 9232267 |
| c33623 | 9317255 |
| c25826 | 9218579 |
| c30164 | 9226137 |
| c25822 | 9217785 |
| c29886 | 9220595 |
| c32314 | 9268194 |
| c24681 | 4035747 |
| c33331 | 9289775 |
| c30141 | 9226742 |
| c26283 | 9230064 |
| c24969 | 9193557 |
| c32339 | 9270758 |
| c31000 | 9244148 |
| c25173 | 9203272 |
| c25237 | 9203845 |
| c30840 | 9239850 |
| c30882 | 9239370 |
| c29623 | 9195413 |
| c28359 | 9278979 |
| c30872 | 9239357 |
| c26167 | 9225879 |
| c32103 | 9265563 |
| c33329 | 9293951 |
| c28262 | 9277579 |
| c27280 | 9253048 |

### Table T2: Primary assays (%-inhibition values)

| Drug Query | Compound ID | PDSP ID | SERT | NET | 5-HT2A | 5-HT2C | |
|---|---|---|---|---|---|---|---|
| Fluoxetine | c29237 | 57994 | -28.79 | -8.73 | 10.26 | -1.78 | |
| | c26692 | 57995 | -30.52 | 21.02 | 3.75 | -5.15 | |
| | c29296 | 57996 | -30.52 | 21.5 | 15.27 | 4.32 | |
| | c25328 | 57997 | -26.76 | 35.82 | 22.15 | 5.24 | |
| | c26732 | 57998 | -15.64 | -2.57 | -1.92 | -1.18 | |
| Paroxetine | c33446 | 57999 | -25.87 | 11.05 | 12.5 | 5.69 | |
| | c33448 | 58000 | -20.59 | 1.69 | 3.75 | -1.08 | |
| | c29676 | 58001 | -24.45 | -6.15 | 24.79 | 10.4 | |
| | c33858 | 58002 | -28.43 | -16.62 | 8.26 | 20.12 | |
| | c29440 | 58003 | -22.69 | -6.55 | 8.63 | 1.72 | |
| | | | D2 | D3 | 5-HT2A | 5-HT2C | |
| Lisuride | c32656 | 58004 | 15.77 | -3.78 | 45.23 | -6.1 | |
| | c26429 | 58005 | 17.55 | 5.08 | -6.43 | 2.32 | |
| | c33965 | 58006 | 15.9 | -1.07 | 1.75 | 4.64 | |
| | c33151 | 58007 | 26.43 | 3.85 | -4.56 | -0.6 | |
| | c31211 | 58008 | 23.13 | -0.78 | -14.94 | -8.81 | |
| (±) PD 128,907 HCl | c25518 | 58009 | 8.46 | -2.37 | -9.32 | -1.02 | |
| | c27884 | 58010 | 19.52 | 8 | 1.59 | -9.82 | |
| | c25477 | 58011 | 18.82 | 12.67 | -3.42 | -9.89 | |
| | c24594 | 58012 | 11.29 | -8.25 | -10 | -6.8 | |
| | c31386 | 58013 | 17.17 | 0.8 | -3.94 | -0.61 | |
| | | | 5-HT2A | 5-HT2B | 5-HT2C | D1 | D2 |
| Chlorpromazine | c34037 | 58014 | 8.75 | 33.64 | 31.01 | 28.4 | 18.35 |
| | c33553 | 58015 | 12.77 | 67.79 | 41.05 | 17.82 | 7.66 |
| | c33255 | 58016 | 25.9 | 48.66 | 26.7 | 12.5 | 6.64 |
| | c31508 | 58017 | 3.85 | 3.23 | -19.39 | 0.09 | 7.31 |
| | c31447 | 58018 | 61.94 | 86.4 | 59.09 | 25.91 | 39.24 |
| Mianserine | c28263 | 58019 | 6.74 | -6.2 | 7.12 | -17.1 | 6.22 |
| | c33507 | 58020 | 28.46 | 75.25 | 3.16 | -3.69 | 21.02 |

| | | | | | | | |
|---|---|---|---|---|---|---|---|
| | c26398 | 58021 | 6.07 | 13.94 | 5.57 | -19.09 | 25.88 |
| | c26521 | 58022 | -7.87 | -2.8 | 5.87 | -3.8 | 2.37 |
| | c33914 | 58023 | -2.97 | 24.18 | 2.75 | -6.73 | 15.17 |
| Loxapine | c30977 | 58024 | 7.2 | 50.77 | 6.67 | -0.42 | 28.22 |
| | c28263 | 58019 | 6.74 | -6.2 | 7.12 | -17.1 | 6.22 |
| | c27284 | 58025 | -8.12 | -10.13 | -4.12 | -3.74 | 14.83 |
| | c33507 | 58020 | 28.46 | 75.25 | 3.16 | -3.69 | 21.02 |
| | c26398 | 58021 | 6.07 | 13.94 | 5.57 | -19.09 | 25.88 |
| Clomipramine | c33623 | 58026 | 78.39 | 92.17 | 82.11 | 40.26 | 67.55 |
| | c25826 | 58027 | -4.81 | 29.57 | 1.24 | -7.56 | 24.41 |
| | c30164 | 58028 | 6.7 | 27.58 | -4.62 | -5.63 | 16.78 |
| | c25822 | 58029 | 0.92 | 0.58 | -8.87 | -4.85 | 14.71 |
| | c29886 | 58030 | 7.41 | 51.77 | 13.86 | 9.62 | 20.43 |

| | | | 5-HT1A | 5-HT7A | D2 |
|---|---|---|---|---|---|
| 8-Hydroxy-DPAT | c32314 | 58031 | 4.54 | 40.03 | 16.15 |
| | c24681 | 58032 | 63.5 | 40.97 | 16.91 |
| | c33331 | 58033 | 11.69 | 51.18 | 19.7 |
| | c30141 | 58034 | -2.36 | 39.83 | 1.06 |
| | c26283 | 58035 | -4.38 | 28.38 | 38.05 |
| | | | H3 | H4 | |
| IMETIT 2HBr | c24969 | 58036 | 8.14 | 12.44 | |
| | c32339 | 58037 | 50.2 | 38.18 | |
| | c31000 | 58038 | 68.59 | 63.29 | |
| | c25173 | 58039 | 33.04 | 6.94 | |
| | c25237 | 58040 | 86.54 | 9.11 | |
| | | | Beta1 | Beta2 | Beta3 |
| Isoproteronol | c30840 | 58041 | 10.83 | #N/A | #N/A |
| | c30882 | 58042 | 12.96 | #N/A | #N/A |
| | c29623 | 58043 | 21.4 | #N/A | #N/A |
| | c28359 | 58044 | 17.81 | #N/A | #N/A |
| | c30872 | 58045 | -0.3 | #N/A | #N/A |
| | | | Alpha1A | Alpha2C | |

| | | | | |
|---|---|---|---|---|
| Corynanthine | c26167 | 58046 | #N/A | 17.78 |
| | c32103 | 58047 | #N/A | 17.16 |
| | c33329 | 58048 | #N/A | 41.07 |
| | c28262 | 58049 | #N/A | 63.58 |
| | c27280 | 58050 | #N/A | 38.2 |

### Table T3: Known-drug pairs related by phenotypic distance but with dissimilar structures

| namex | namey | twin-nn dist | tanimoto dist | correlation dist | novelty | ChEMBL_Similarity |
|---|---|---|---|---|---|---|
| 7-Hydroxy-DPAT HBr | Ropinirole HCl | 0.254 | 0.543 | 0.269 | 1.288 | 0.636 |
| (±)-PD 128,907 HCl | Ropinirole HCl | 0.233 | 0.567 | 0.275 | 1.335 | 0.556 |
| (+)-PD 128907 | Ropinirole HCl | 0.224 | 0.567 | 0.277 | 1.343 | 0.556 |
| trans-7-Hydroxy-PIPAT maleate | LY-163,502 2HCl | 0.136 | 0.505 | 0.211 | 1.369 | 0.500 |
| 1-[1-(2-Benzo[b]thienyl)cyclohexyl)]piperidine maleate | PRE-084·HCl | 0.268 | 0.574 | 0.234 | 1.306 | 0.500 |
| Ropinirole HCl | Lisuride maleate | 0.132 | 0.549 | 0.323 | 1.417 | 0.444 |
| 7-Hydroxy-DPAT HBr | Pergolide mesylate | 0.257 | 0.576 | 0.168 | 1.319 | 0.438 |
| Ropinirole HCl | trans-7-Hydroxy-PIPAT maleate | 0.092 | 0.544 | 0.280 | 1.453 | 0.400 |
| LY-163,502 2HCl | (±)-SKF-82958 HBr | 0.228 | 0.534 | 0.229 | 1.306 | 0.400 |
| LY-163,502 2HCl | R(+)-6-BROMO-APB HBr | 0.192 | 0.529 | 0.326 | 1.337 | 0.400 |
| Mianserin HCl | Oxotremorine sesuifumarate | 0.285 | 0.833 | 0.164 | 1.547 | 0.378 |
| Ropinirole HCl | LY-163,502 2HCl | 0.103 | 0.578 | 0.249 | 1.475 | 0.375 |
| trans-7-Hydroxy-PIPAT maleate | Lisuride maleate | 0.146 | 0.514 | 0.175 | 1.367 | 0.375 |
| Yohimbine HCl | Oxymetazoline HCl | 0.147 | 0.548 | 0.150 | 1.401 | 0.375 |
| 7-Hydroxy-DPAT HBr | Lisuride maleate | 0.254 | 0.579 | 0.191 | 1.325 | 0.364 |
| Clozapine | Chlorpromazine HCl | 0.227 | 0.546 | 0.248 | 1.319 | 0.337 |
| Clozapine | Chlorpromazine HCl | 0.215 | 0.546 | 0.214 | 1.331 | 0.337 |
| 7-Hydroxy-DPAT HBr | Bromocriptine mesylate | 0.187 | 0.594 | 0.323 | 1.408 | 0.333 |
| 7-Hydroxy-DPAT HBr | LY-163,502 2HCl | 0.249 | 0.534 | 0.173 | 1.285 | 0.300 |
| trans-7-Hydroxy-PIPAT maleate | R(-)-Propylnorapomorphine HCl | 0.277 | 0.523 | 0.350 | 1.246 | 0.300 |
| Mianserin HCl | Chlorpromazine HCl | 0.199 | 0.538 | 0.136 | 1.339 | 0.288 |
| Bromocriptine mesylate | Ropinirole HCl | 0.155 | 0.538 | 0.340 | 1.382 | 0.286 |
| Mianserin HCl | WB 4101 HCl | 0.228 | 0.714 | 0.170 | 1.486 | 0.283 |
| Chlorpromazine HCl | (+)-Butaclamol HCl | 0.147 | 0.530 | 0.181 | 1.383 | 0.279 |
| Chlorpromazine HCl | Pergolide mesylate | 0.217 | 0.554 | 0.222 | 1.337 | 0.268 |

| | | | | | | |
|---|---|---|---|---|---|---|
| trans-7-Hydroxy-PIPAT maleate | Pergolide mesylate | 0.290 | 0.542 | 0.190 | 1.252 | 0.267 |
| Clothiapine | Thioridazine HCl | 0.287 | 0.502 | 0.290 | 1.215 | 0.263 |
| cis-(±)-N-methyl-N-[2-(3,4-dichlorophenyl)ETHYL]-2-(1-pyrrolidinyl)cyclohexamine | 1-[1-(2-Benzo[b]thienyl)cyclohexyl)]piperidine maleate | 0.192 | 0.610 | 0.163 | 1.419 | 0.250 |
| cis-(±)-N-methyl-N-[2-(3,4-dichlorophenyl)ETHYL]-2-(1-pyrrolidinyl)cyclohexamine | PRE-084·HCl | 0.242 | 0.687 | 0.206 | 1.445 | 0.250 |
| 6-Nitroquipazine maleate | Bopindolol malonate | 0.140 | 0.603 | 0.261 | 1.463 | 0.250 |
| PNU 96415E | trans-7-Hydroxy-PIPAT maleate | 0.232 | 0.560 | 0.318 | 1.328 | 0.250 |
| B-HT 920 2HCl | LY-163,502 2HCl | 0.149 | 0.585 | 0.160 | 1.435 | 0.250 |
| Ropinirole HCl | R(-)-Propylnorapomorphine HCl | 0.256 | 0.538 | 0.380 | 1.282 | 0.250 |
| trans-7-Hydroxy-PIPAT maleate | (±)-SKF-82958 HBr | 0.257 | 0.514 | 0.228 | 1.257 | 0.250 |
| trans-7-Hydroxy-PIPAT maleate | R(+)-6-BROMO-APB HBr | 0.237 | 0.520 | 0.311 | 1.282 | 0.250 |
| RS 17053 HCl | Naftopidil HCl | 0.071 | 0.564 | 0.139 | 1.493 | 0.250 |
| Clozapine | WB 4101 HCl | 0.173 | 0.704 | 0.224 | 1.531 | 0.241 |
| Clozapine | WB 4101 HCl | 0.157 | 0.704 | 0.179 | 1.547 | 0.241 |
| PNU 96415E | WB 4101 HCl | 0.280 | 0.710 | 0.272 | 1.430 | 0.240 |
| Cyproheptadine HCl | Mianserin HCl | 0.280 | 0.559 | 0.390 | 1.279 | 0.239 |
| Mianserin HCl | Clomipramine HCl | 0.287 | 0.530 | 0.147 | 1.243 | 0.239 |
| Bromocriptine mesylate | trans-7-Hydroxy-PIPAT maleate | 0.167 | 0.534 | 0.336 | 1.366 | 0.231 |
| Thioridazine HCl | Imipramine HCl | 0.069 | 0.527 | 0.198 | 1.458 | 0.231 |
| 6-Nitroquipazine maleate | Nisoxetine HCl | 0.175 | 0.702 | 0.180 | 1.527 | 0.222 |
| PNU 96415E | B-HT 920 2HCl | 0.155 | 0.658 | 0.266 | 1.503 | 0.222 |
| SCH 23390 HCl | AMI-193 | 0.206 | 0.588 | 0.208 | 1.382 | 0.222 |
| Mianserin HCl | Thioridazine HCl | 0.293 | 0.505 | 0.205 | 1.212 | 0.217 |
| WB 4101 HCl | RS 17053 HCl | 0.258 | 0.603 | 0.261 | 1.346 | 0.217 |
| Chlorpromazine HCl | Imipramine HCl | 0.163 | 0.520 | 0.154 | 1.357 | 0.214 |
| Loxapine succinate | Chlorpromazine HCl | 0.198 | 0.559 | 0.277 | 1.360 | 0.209 |
| Chlorpromazine HCl | Methiothepin maleate | 0.094 | 0.559 | 0.413 | 1.465 | 0.209 |
